# Aquamin: A Source of Bioavailable Calcium, Magnesium and Additional Cationic Trace Elements in Colon Health and Beyond

**DOI:** 10.64898/2026.10.14.759236

**Authors:** Shannon D McClintock, Muhammad N Aslam, Randall N Knibbs, Jacob E Theisen, Keith MacRenaris, Shane O’Connell, Denise M O’Gorman, James Varani

## Abstract

**Introduction:** Aquamin is a calcium– and magnesium-rich natural product derived from marine red algae that contains multiple trace elements. Human and mouse studies have demonstrated potential health benefits.

**Methods:** Aquamin and calcium carbonate were compared using a two-stage simulated gastric and intestinal digestion protocol. ICP-OES/ICP-MS was used to quantify cationic elements in soluble and insoluble fractions. Binding of dispersed mineral powders to human colon epithelial cells was assessed by light and scanning electron microscopy, and uptake was assessed by transmission electron microscopy. TMT-based proteomics identified protein changes in colon epithelial organoids following mineral exposure. Serum samples from 90– and 180-day Aquamin intervention trials were analyzed for changes in circulating cationic elements.

**Results:** Both Aquamin and calcium carbonate were solubilized under simulated gastric conditions, although Aquamin was more acid-resistant. After transfer to neutral-pH intestinal conditions, limited precipitation occurred, and most calcium remained soluble. With Aquamin, magnesium and strontium also remained soluble, whereas several trace elements became partially or completely insoluble. Aquamin rapidly bound to colon epithelial cells, with ultrafine Aquamin particles detected between adjacent cells. Calcium carbonate bound more slowly and to fewer cells. Aquamin increased proteins involved in enterocyte cation uptake, intracellular trafficking, and basolateral secretion. Serum strontium levels increased in study participants after 90 or 180 days of Aquamin consumption; most other elements changed minimally, although serum iron increased modestly.

**Conclusions:** Aquamin provides a bioavailable source of several cationic elements. Soluble calcium, magnesium, and strontium may be absorbed primarily through transcellular pathways, while less-soluble elements may also cross the colonic epithelium through paracellular mechanisms.

## INTRODUCTION

An adequate daily intake of calcium is essential for health because this element is a critical component of virtually every biological process (Bootman and Bultynck, 2020; Stewart et al. 2015). Studies carried out over the past fifty years have repeatedly demonstrated that a majority of individuals in Western countries do not achieve the recommended daily intake of this important element (Peterlik et al. 2009; Heaney 2003). Deficiencies in other important trace elements have also been documented (Camaschella 2019; Rolić et al. 2025). The Western style diet, with its reliance on highly-processed food, is largely responsible for mineral deficiencies in advanced-economy countries. In the west, trace element deficiencies tend to be chronic, sub-clinical and population group-specific. Mineral deficiencies are also prevalent globally (López-Alonso et al. 2025; Passarelli et al. 2022; Stevens et al. 2022). In developing countries and regions where intensive farming is widespread, especially where the diet is primarily plant-based, mineral deficiencies often result from nutrient/mineral-depleted soils. Here mineral deficiencies are often associated with overt diseases or with compromise of anti-disease mechanisms. While “healthy eating” is always the preferred option for achieving a balanced intake of essential trace elements, the use of mineral supplements could be part of the strategy for preventing deficiencies.

Aquamin is a calcium-rich, magnesium-rich, multi-trace element natural product obtained from the skeletal remains of marine red algae (Adey and McKibben 1970). Our own long-term (15-18 month) studies in mice have documented substantial health benefits when Aquamin was included in the diet over the life of the animal. Benefits include reduced tumor formation throughout the gastrointestinal tract (Aslam et al. 2010a; Aslam et al. 2012a) and in the liver (Aslam et al. 2012b), decreased ulcerative dermatitis (Hampton et al. 2015) and increased bone mineral content and bone strength (Aslam et al. 2010b; Aslam et al. 2013; Aslam et al. 2016). In these studies, a substantial lessening of inflammation throughout the gastrointestinal tract and systemically was seen in conjunction with health benefits. In support of an anti-inflammatory mechanism, studies by Aviello et al. (2014) showed that including Aquamin in the drinking water of IL-10 −/− mice reduced the incidence and severity of colitis.

Interventional trials in humans suggest that health benefits of Aquamin extend to people. In a 90-day study with healthy individuals, daily Aquamin ingestion was associated with a change in the bacterial population of the gut and with reduced levels of several bile acids and increased levels of short chain fatty acids (Aslam et al. 2020a). A proteomic screen identified increased expression of several proteins that contribute to epithelial cohesion and barrier integrity (Aslam et al. 2021). In a follow-up 180-day study conducted with subjects having mild ulcerative colitis (UC) or in-remission disease, improvement in several disease-related biomarkers was observed (Aslam et al. 2025a) along with increased expression of the same barrier proteins. A subsequent study in UC subjects (Aslam et al. 2026) provided direct evidence for improved barrier function *in vivo*. Consistent with these findings, improved barrier protein expression was observed in human colon organoid cultures (Attili et al. 2019; Aslam et al. 2020b; McClintock et al. 2020; Varani et al. 2022; Varani et al. 2023; Aslam et al. 2025b). In the *in vitro* studies, increased barrier protein expression was accompanied by improved organoid cohesion and by increased trans-epithelial electrical resistance (McClintock et al. 2020; Varani et al. 2022; Varani et al. 2023).

In both the preclinical studies and the interventional trials, calcium alone at concentrations comparable to amounts delivered in Aquamin also demonstrated bioactivity but did not provide the same changes as seen with Aquamin, nor the same level of benefit in some cases. Functional differences between Aquamin and calcium alone may reflect the combined beneficial activities of the additional trace elements provided along with calcium in the algae mineral preparation. This would not be surprising because several of the elements present in Aquamin are well known to be required for health (Camaschella 2019; Rolić et al. 2025; López-Alonso et al. 2025; Liu and Dudley 2025; Rosanoff et al. 2012; Prasad 2013; Roohani et al. 2013; Duncan et al. 2015; Melkote et al. 2026; Wang et al. 2023; Rayman 2012; Cardoso et al. 2015; Jiang et al. 2018; Chen et al. 2020; Boivin et al. 1996; Marie et al. 2001).

Alternatively, the beneficial effects of Aquamin may be due, primarily, to calcium (the most abundant cationic element in the algae preparation and a critical component of virtually every biological process [Bootman and Bultynck 2020; Stewart et al. 2015; Peterlik et al. 2009; Heaney 2003]) but reflect differences between Aquamin and calcium alone in bioavailability. Specifically, the multi-trace element product may differ from a pure calcium salt in properties that affect acid-buffering capacity, or how the salt undergoes solubilization in the acid environment of the stomach and/or re-precipitation in the higher pH environment of the small intestine. In addition, structural differences between the two materials could influence how insoluble components interact with cells lining the gastrointestinal wall. The studies described in this report begin to address these issues. Here we have assessed Aquamin and calcium carbonate solubilization and re-precipitation in a two-stage gastric– and intestinal-simulating digestion protocol (Minekus et al. 2014; Felice et al. 2018). In parallel, Aquamin and calcium salts were compared for ability to bind to and interact with colonic epithelial cells. Finally, as a way to confirm trace element delivery by Aquamin in humans, we compared trace element levels in serum samples obtained before and after intervention with Aquamin in two recent clinical trials (Aslam et al. 2020a; Aslam et al. 2021; Aslam et al. 2025a).

## MATERIALS AND METHODS

### Aquamin

Aquamin is a calcium-rich, magnesium-rich, multiple trace element natural product obtained from the skeletal remains of red marine algae of the *Lithothamnion* family (Adey and McKibben 1970) (Marigot Ltd, Cork, Ireland). The US Food and Drug Administration (FDA) has approved Aquamin as a GRAS-listed product (GRAS 000028), and it is used as a food supplement for human consumption in Europe, Asia, Australia, and North America. Aquamin is also used as an agricultural supplement.

Aquamin exists as an aqueous-insoluble powder at neutral pH. When dispersed, particle sizes range from less than 1 μm to approximately 30 μm in diameter. Aquamin has a calcium concentration of approximately 32% and a magnesium concentration of 2.6%. Detectable amounts of up to 72 trace elements can be identified in the algal product. Among these are several nutritionally important cationic elements including strontium, iron, manganese, copper, zinc, chromium, and selenium. The cationic elements in Aquamin are, primarily, in the form of carbonate salts (calcite, vaterite and aragonite). The material contains organic matter at approximately 3.4% w/w, which is not present in calcium carbonate sources. The nature of the organic matter is predominately sulphated carbohydrate, but other molecules may also exist. Supplement Table 1 provides a list of 12 important cationic elements detected in Aquamin and their relative abundance. The preparation used here is referred to as Aquamin-F; this is the form that has been used in the past clinical trials conducted in our laboratory (Aslam et al. 2020a; Aslam et al. 2021; Aslam et al. 2025a; Aslam et al. 2026). For past *in vitro* studies, we used a soluble-fraction of Aquamin, known as Aquamin-Soluble (Attili et al. 2019; Aslam et al. 2020b; McClintock et al. 2020; Varani et al. 2022; Varani et al. 2023; Aslam et al. 2025b).

### Calcium carbonate

Calcium carbonate was used as a control. The insoluble salt was prepared by suspending 200 μL of a 0.5 M solution of calcium chloride with excess sodium dibasic carbonate. The precipitate that formed was centrifuged, washed ten times with double-distilled water and then dried. The dried material was pulverized with mortar and pestle at the time of use. In initial experiments, commercially available sources of calcium for human and animal feed application were obtained from commercial sources. These included limestone flour, ground oyster shell, ground chicken eggshell and monocalcium phosphate salt.

### Two-stage digestion simulation protocol

Simulation of *in vivo* digestion of Aquamin and calcium carbonate was carried out following the procedure of Minekus et al. (2014) and Felice et al. (2018). Stage-1 (simulated gastric stage) consisted of an inorganic aqueous salt buffer containing the enzyme pepsin (2000 U/mL). Both materials were added at 800 mg in 20 mL of stage-1 buffer. The ratio of Aquamin to stage-1 buffer was chosen to “meet or exceed” the expected conditions of a research participant in our recent clinical studies (Aslam et al. 2020a; Aslam et al. 2021; Aslam et al. 2025a; Aslam et al. 2026). In clinical studies, participants consumed two 600-mg Aquamin capsules at a time (containing 400 mg of calcium in total); the stomach volume was assumed to be greater than 40 mL (Schubert and Kaunitz 2021). Then, 6M hydrochloric acid (HCl) was added dropwise to bring the pH to 3.0. The simulated gastric fluid was incubated at 37°C with stirring for two hours. The pH was checked after one hour but no additional acid was needed to maintain the pH at 3. At the completion of the gastric simulation (first stage), two 1-mL samples were collected. One was saved as is; the other was passed through a 0.22 μm polycarbonate filter. Both non-filtered and filtered fractions were analyzed for levels of calcium, magnesium, strontium, iron, manganese, copper, zinc, chromium and selenium as well as for three lanthanides – i.e., lanthanum, cerium and gadolinium by ICP-OES (calcium and magnesium) or ICP-MS for all the others (see below).

The remainder of the stage-1 samples were diluted 1:2 with the simulated intestinal fluid (stage-2) buffer. The simulated intestinal fluid consisted of an inorganic salt solution to which pancreatin (1000 U/mL), and bile salts (10 mM) were added. The acidic solution was brought to pH 7 with 1N NaOH. The simulated intestinal fluid was incubated at 37°C with stirring for two hours; additional NaOH was provided as needed after one hour to maintain neutral pH. At the completion of the intestinal simulation, two samples were saved for analysis by ICP-OES or ICP-MS as after stage-1.

Both Aquamin and calcium carbonate samples appeared to be entirely in solution at the end of the (acid pH) stage-1 gastric digestion. However, a visible precipitate formed during the (neutral pH) stage-2 intestinal digestion. Centrifugation (1000xg for 10 minutes) was done to separate the precipitated material, which was then washed two times in calcium– and magnesium-free PBS and analyzed for levels of the same twelve elements. The precipitated materials were also examined by Scanning electron microscopy (SEM; see below).

### Inductively Coupled Plasma (ICP) Analysis

Elemental analysis of samples obtained from digestion studies and of serum samples was performed at the Michigan State University Quantitative Bio Element Analysis and Mapping (QBEAM) Center. Detailed methods for elemental analysis using inductively coupled plasma–optical emission spectrometry (ICP-OES) and triple-quadrupole inductively coupled plasma–mass spectrometry (ICP-QQQ-MS), including sample and standard preparation, instrumentation, operating parameters, analytes, calibration, and quality-control procedures, are provided in Supplementary File 1.

### Scanning electron microscopy (SEM) and transmission electron microscopy (TEM)

SEM was used to visualize Aquamin and calcium carbonate powders and as a way to obtain high-definition views of the powdered materials bound to the cell surface. TEM was used to assess particle localization of the bound Aquamin powder. SEM and TEM analyses were done in conjunction with the Microscopic Imaging Core Laboratory at the University of Michigan. For SEM evaluation of the dispersed powders, EM stubs with an adhesive disc were prepared by lightly dusting the powders over the adhesive surface and then gently tamping them down on a piece of parafilm to ensure the particles were adhered. Cells for SEM were grown on Thermanox membranes (174977; ThermoFisher). Subsequent processing included fixation with 2.5% glutaraldehyde in 0.1M sodium cacodylate buffer and visualization using a Zeiss EVO 15 scanning electron microscope. For TEM, cells were grown in 60 mm dishes to approximately 75% confluence. Cells were scraped and pelleted. Subsequent processing included fixation with 2.5% glutaraldehyde in 0.1M sodium cacodylate buffer followed by post-fixation with Osmium tetroxide, dehydration and embedding in resin which was then polymerized in an oven at 60°C to facilitate thin-sectioning. Cells were visualized using a JEOL JEM-1400 Plus TEM.

### Assessment of Aquamin binding to colonic epithelial cells

Human colon-derived epithelial cells (Caco-2) were obtained from the American Type Culture Collection. Caco-2 is an adherent colon cell line derived from a colorectal adenocarcinoma (HTB-37, ATCC) (Trainer et al. 1988). Cells were grown in monolayer culture using DMEM containing 20% fetal bovine serum as growth medium. Growth was at 37°C in an atmosphere of 95% air and 5% CO_2_. Cells were subcultured as needed. For binding studies, cells were plated in chambers of a 4-chamber glass slide at 2×10^4^ cells per chamber in growth media and allowed to proliferate until 50-75% of the surface area of the chamber was covered. The slide chambers were then gently washed two times in PBS. Aquamin or calcium carbonate was added to the chambers in a 0.5 mL volume at final concentrations between 40 and 200 μg/mL. Mineral binding to the cell surface was assessed microscopically at times between 2 hours and 48 hours. At each time-point, the slides were gently washed two times in PBS followed by formalin-fixation and staining with hematoxylin and eosin. Stained cells were digitized using the Aperio Slide Scanning System. The digitized slides were evaluated by light microscopy. Total cells and cells with visible attached particles were counted, and the percentage of cells with particles attached was calculated. Cells with bound particles were also visualized by SEM and TEM as described above.

Mineral binding to human colon organoid-derived epithelial cells was also assessed. The UM-Human Organoid Core was the source of these cells. The collection and use of human colon tissue for this purpose was approved by the Institutional Review Board (IRBMED) at the University of Michigan and all subjects provided written informed consent prior to biopsy. Organoid derived colon epithelial cells were obtained from growing organoids by dissociating the tissue into small cell aggregates (less than 40 µm in size) and plating onto collagen IV-coated transwells (0.4 µm pore size, 0.33 cm² PET, Corning Costar) at a density of 2×10^5^ aggregates per transwell insert. When nearly complete monolayers had developed, the growth medium used in the core laboratory was replaced with a 25:75 mix of L-WRN and KGM Gold culture media as described in our previous organoid culture studies (Attili et al. 2019; Aslam et al. 2020b; McClintock et al. 2020; Varani et al. 2022; Varani et al. 2023; Aslam et al. 2025b). The final calcium concentration in the medium was 0.25 mM, and the final serum concentration was 2.5%.

### Proteomic assessment

Colon organoid data from 12 subjects and 14 independent proteomic runs, previously analyzed individually across four independent studies and described in detail previously (Attili et al. 2019; Aslam et al. 2020b; Varani et al. 2023; Aslam et al. 2025b), were included here. From each subject, 50 µg of protein from control colon organoid cultures (0.25 mM calcium) or matched cultures treated with a level of Aquamin to bring the final calcium concentration to 3 mM was trypsin-digested and labeled with Tandem Mass Tags. Equal amounts of labeled peptides were then pooled, fractionated by two-dimensional liquid chromatography, and analyzed on an Orbitrap Fusion Tribrid mass spectrometer using MultiNotch MS3 quantification. Spectra were processed in Proteome Discoverer and searched against a human UniProt/SwissProt database. Protein and peptide identifications were limited to a false-discovery rate of ≤2%, with quantification based on high-quality MS3 spectra. Differential abundance was determined after normalizing Aquamin-treated samples to the corresponding control for each subject. Results from individual matched data sets were then merged across all 12 subjects to provide a comprehensive assessment of the response to Aquamin intervention. UniProt annotations were used to identify proteins of interest (i.e., for the present study, these were proteins involved in trace element transport across the gastrointestinal wall). Reactome database (reactome.org) was used for pathway enrichment analyses. Additionally, from a subset of subjects (n=3), Aquamin (providing 3 mM calcium) was compared to calcium alone at the same level for the same set of proteins (see Attili et al. 2019). The mass spectrometry proteomics data set from each study was deposited to the ProteomeXchange Consortium via the PRIDE partner repository for open access.

### Cationic element assessment in human serum samples

Cationic element levels were assessed in serum samples obtained from participants in two recently-completed Aquamin interventional trials. The first trial was conducted with healthy subjects [ClinicalTrials.gov ID NCT02647671] (Aslam et al. 2020a; Aslam et al. 2021) and the second with in-remission UC patients or those with mild disease [ClinicalTrials.gov ID NCT03869905] (Aslam et al. 2025a). These investigator-initiated interventional trials were conducted with Aquamin under an FDA-approved IND. Oversight was provided by the Institutional Review Board at the University of Michigan Medical School (IRBMED: HUM00076276 and HUM156676). In both trials, separate cohorts consumed two 600-mg Aquamin capsules twice per day for either 90– or 180-days (providing 800 mg of calcium per day). In both studies, additional subjects consumed maltodextrin for 90 days as a placebo. Prior to intervention and at day-90 and day-180, blood was drawn as part of a safety evaluation and for assessment of clinical and mechanistic biomarkers. Remaining serum was saved and eventually used for elemental analysis in the current study. For this, serum samples were diluted 1:100 in 3% Nitric acid and subjected to ICP-OES and ICP-MS as described above.

### Statistical analysis

Group means and standard deviations were calculated for discrete datasets. The resulting data were analyzed using analysis of variance (ANOVA), followed by a two-tailed paired (for pre-post analysis) and unpaired t-test for group comparisons in GraphPad Prism version 11. Proteomic data were initially processed in Proteome Discoverer, which calculated abundance ratios and *p* values using a nested design incorporating biological replicates. Sample-group *p* values were determined using Tukey’s HSD post hoc test following ANOVA. Statistical significance was defined as *p* < 0.05.

## RESULTS

### Aquamin solubilization and re-precipitation in the two-stage digestion simulation

In the first set of studies, 100 mg of Aquamin was suspended in 2 mL of saline and exposed to 1 M HCl in increasing amount to determine how much acid would be required to bring the solution to pH 3.5 and completely solubilize the mineral product. Similar amounts of calcium salts from various sources (mostly carbonate) were evaluated in comparison. Results presented in Supplement Figure 1 show that the total volume of acid needed for the aqueous-insoluble material to reach pH 3.5 was greater with Aquamin than with calcium from any of the sources. The time to reach pH 3.5 was correspondingly longer with Aquamin than with the calcium salts.

Based on the results of this preliminary study, 800 mg of Aquamin or calcium carbonate was brought up in 20 mL of the simulated gastric fluid and the complete stage-1 and stage-2 digestion simulation protocol carried out in sequence as described in the Materials and Method Section. At the completion of each stage of the digestion protocol, samples were collected and analyzed. As shown in Figure 1A, virtually all of the initially present calcium in Aquamin was solubilized (i.e., defined as filterable through a 0.22 μm filter) in the stage-1 (gastric simulation). Most of the calcium was still soluble at the conclusion of the (intestinal simulation) stage-2. Similar results were obtained with magnesium and strontium. That is, both solubilized in the gastric-simulating stage and remained soluble in the intestine-simulating stage. Figure 1A also provides results for nine additional trace elements. At the completion of gastric stage-1, the soluble form (i.e., filterable through a 0.22 μm filter) predominated with all nine elements. With the exception of iron, copper, and chromium, >99% solubility was achieved. With iron, 69% was soluble after the gastric-digestion simulation; with copper, 83% was soluble and with chromium 97% was in solution.

**Figure 1.**
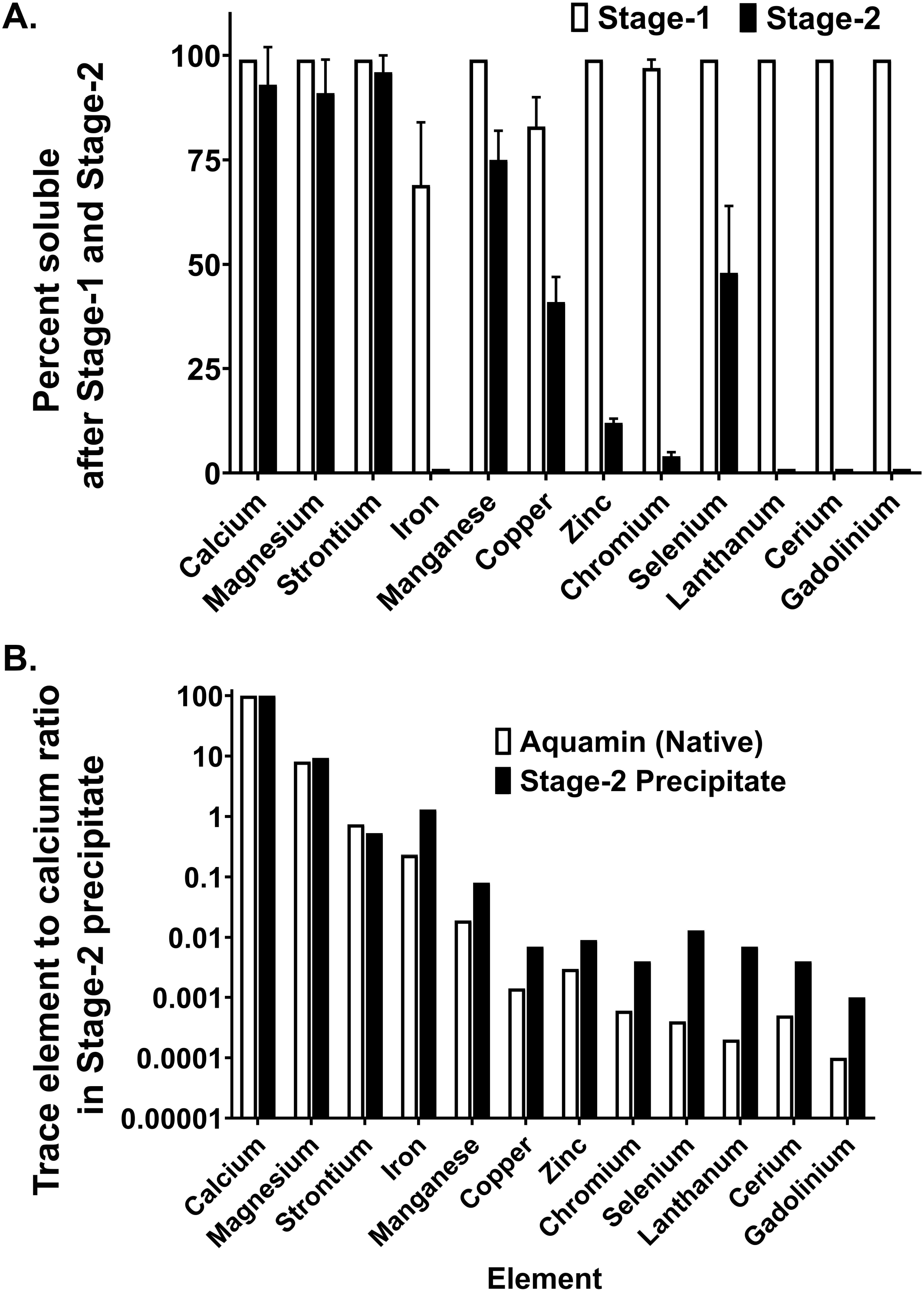
Solubilization and re-precipitation of Aquamin in the two-stage digestion simulation. **A**. Distribution of cationic elements in digestion buffers at the end of stage-1 and stage-2. Samples collected at the end of stage-1 were assayed “as is”. Samples collected at the end of stage-2 were centrifuged to remove the precipitate that had formed during incubation and then assayed. Prior to assay, a portion of each sample from both stages was passed through a 0.22 μm polycarbonate filter. Then, levels of each element of interest in both the non-filtered and filtered samples were determined by ICP-OES (calcium and magnesium) or by ICP-MS (strontium, iron, manganese, copper, zinc, chromium, selenium, lanthanum, cerium, and gadolinium). Levels of each element were compared in filtered and unfiltered samples. Values represent means and standard deviations based on four separate experiments. **B**. Distribution of cationic elements in the precipitate that formed during Stage-2. Levels of each element of interest were determined by ICP-OES (calcium and magnesium) or by ICP-MS (strontium, iron, manganese, copper, zinc, chromium, selenium, lanthanum, cerium, and gadolinium). The calcium value was set at 1.0 and the value of other elements expressed as a ratio to calcium. The ratio of each element to calcium in native Aquamin was assessed in parallel for control. Values represent means and standard deviations based on three separate experiments.

At the conclusion of the intestinal simulation (stage-2), approximately 80-85% (by weight) of initial Aquamin was soluble, while the recovered precipitated material represented the remaining 15-20%. At that stage, the distribution between soluble and insoluble fractions was different for each of the elements (Figure 1A). Most of the calcium, magnesium and strontium remained in the soluble fraction at the end of the intestinal simulation. Manganese was 75% soluble post-stage-2, whereas copper, zinc, chromium, and selenium were 41%, 12%, 4%, and 48% soluble, respectively. With iron, less than 1% of the material was recovered in the soluble fraction. Similarly, while all three lanthanides were soluble at the completion of gastric (stage-1) digestion, all were almost completely insoluble at the end of intestinal stage-2 (Figure 1A).

Although the majority of the solubility data were obtained at the completion of the two-hour incubation period in the stage-1 and stage-2 buffers, a final experiment was conducted in which samples were taken after 5, 10, 30, 60, 90 and 120 minutes of incubation in the neutral pH, intestinal-mimicking stage-2 buffer. The results presented in Supplement Table 2 demonstrate that equilibrium was reached rapidly with each element after transferring to the second stage buffer. That is, solubility data at the 5-minute collection point were not substantially different from the data obtained at the end of the two-hour incubation with any of the 12 elements.

As part of the analysis, the mineral content of the visible precipitate formed during the intestinal stage-2 simulation was also evaluated separately. Figure 1B shows the level of each element in the stage-2 precipitate relative to the level of calcium. For comparison, the level of each element relative to calcium in intact Aquamin is also shown. As expected, the magnesium-to-calcium and strontium-to-calcium ratios in the precipitate were similar to the corresponding ratios to calcium in intact Aquamin. In contrast, the relative levels of the other nine cationic elements were higher in the intestinal stage-2 precipitate than in intact (predigested) Aquamin. On a per-weight basis, iron content was increased the most.

Along with elemental analysis, we examined the stage-2 precipitate by SEM. Both intact Aquamin and intact calcium carbonate were assessed for comparison. As shown in Supplement Figure 2, native Aquamin particles showed a wide range of size (<1 μm to greater than 35 μm in cross section) while the calcium carbonate particles had a narrow size range (30-40 μm). The stage-2 precipitate was indistinguishable from intact Aquamin.

Finally, calcium carbonate was examined in the two-stage digestion protocol in parallel with Aquamin. As seen with Aquamin, the calcium carbonate suspension appeared (visually) to have undergone complete solubilization in gastric stage-1 and also like Aquamin, partially reprecipitated during intestinal stage-2. As expected, most of the calcium was in the soluble form at the end of both stage-1 (98±1% soluble) and stage-2 (98±2% soluble). Also, as expected, there was essentially no significant amount of any other cationic element in fractions derived from calcium carbonate after either stage. Values obtained with calcium carbonate are presented in Supplement Table 3.

### Aquamin binding to colon epithelial cells

Caco-2 cells have been extensively utilized to study binding and internalization of three-dimensional particles (Perfecto et al. 2017; Pereira et al. 2013; Domenech et al. 2021; Cubadda et al. 2026). These cells were, therefore, used in our studies to assess Aquamin binding. The quantitative data shown in Figure 2A demonstrate that Aquamin binding to the epithelial cell surface occurred within two hours of exposure, and that the percentage of cells with bound Aquamin increased over the 48-hour incubation period. Figure 2A also demonstrates calcium carbonate binding. Fewer cells had detectable material bound at each time point.

**Figure 2.**
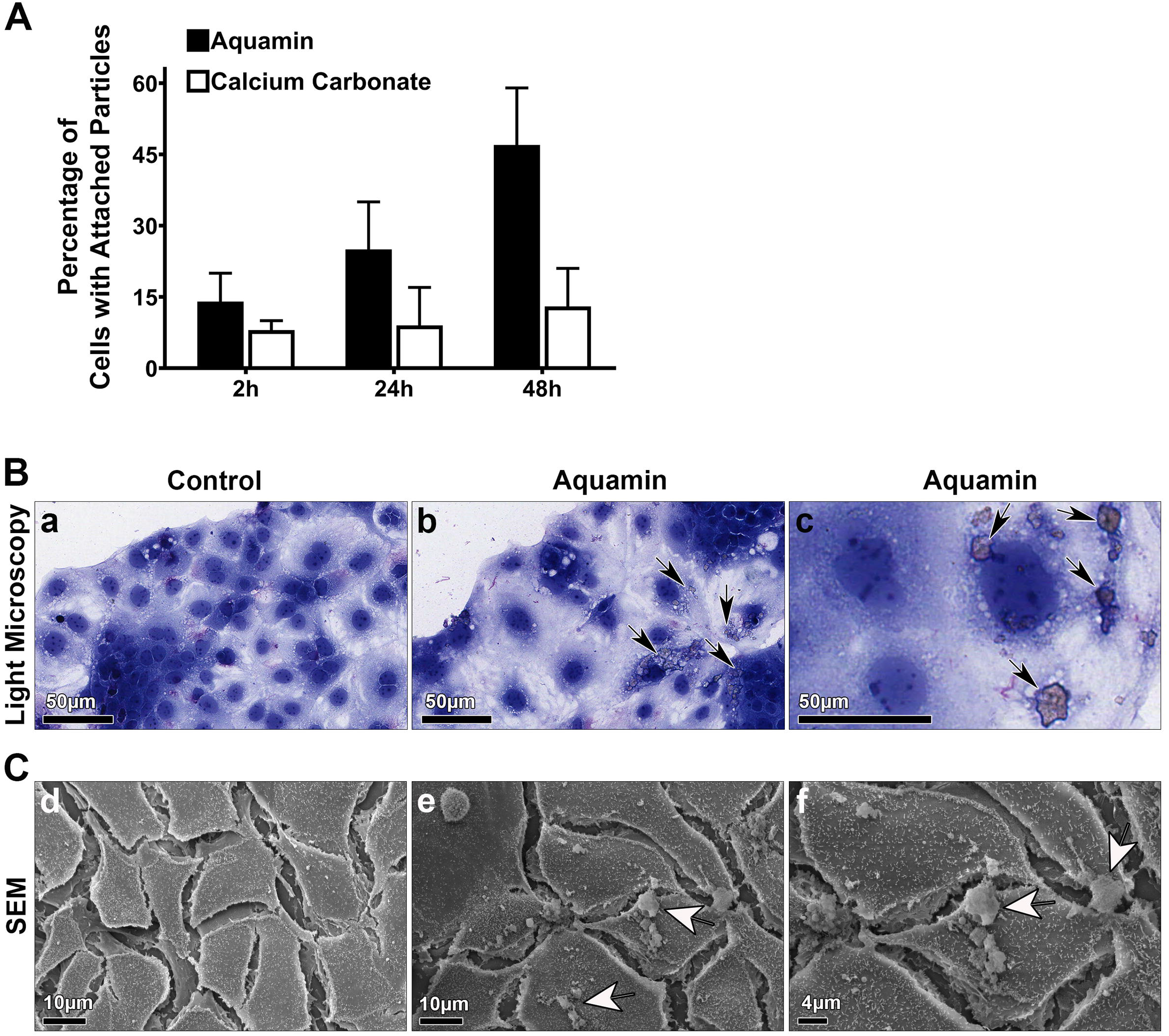
Aquamin binding to Caco-2 cells in monolayer culture. **A**. Quantitation of binding: Caco-2 cells were grown on 4-chamber glass slides and binding studies with Aquamin and calcium carbonate were carried out as described in the Materials and Methods Section. Values shown represent the percentage of cells with visible particles on the cell surface as determined from light microscopic images. Values are means and standard deviations based on ten 400X fields per data point in one experiment. The experiment was carried out three times with similar results. **B**. Light microscopy: control Caco-2 cells and Aquamin-treated Caco-2 cells. Arrows point to bound Aquamin particles. Bars in panels a, b, and c = 50 μm. **C**. Scanning electron microscopy (SEM): control Caco-2 cells and Aquamin-treated Caco-2 cells. Arrows point to bound Aquamin particles. Bars in panel d and e = 10 μm and bar in panel f = 4 μm. Panels a, b, d and e present lower magnification and panels c and f present higher magnification images.

Figure 2B presents light microscopic images of control Caco-2 cells and cells exposed to Aquamin. As seen in the light microscopic images, control Caco-2 cell monolayer (panel a) consisted of mostly small, closely-packed cells with dark-staining nuclei and visibly light-stained cytoplasm. A high cytoplasm-to-nuclear ratio (reflecting differentiation) characterized these cells. Aquamin-exposed cells appeared similar to those in the control panel, except that bound particles were evident with many of the cells (arrows in panels b and c).

Assessment at the SEM level is depicted in Figure 2C (panels d-f). The control panel (d) revealed a monolayer of flat epithelioid cells with numerous tiny surface projections (microvilli). Except for the occasional dust particles, virtually no bound material was evident. When Aquamin-exposed cells were examined, bound particles could be seen on the cell surface. Panel e is a low power image showing several cells with bound particles. The higher-magnification image (panel f) depicts Aquamin binding at an area where contact between two cells is evident.

Figure 3 presents results from TEM evaluation. Panel a demonstrates ultrastructural features observed in a typical control Caco-2 cell. White arrows in the panel delineate the boundary between two adjacent cells. In panels b and c, images of Caco-2 cells following exposure to Aquamin for 48 hours are presented. The Aquamin-exposed cells shown in the image are similar in appearance to control cells. The presence of submicron-sized electron-dense particles in the intercellular space between two adjacent cells is seen (black arrows). Panel c shows a higher magnification view of the boxed area in panel b, highlighting the electron-dense particles (60-100 nm in cross section) in the intercellular space between the two cells.

**Figure 3.**
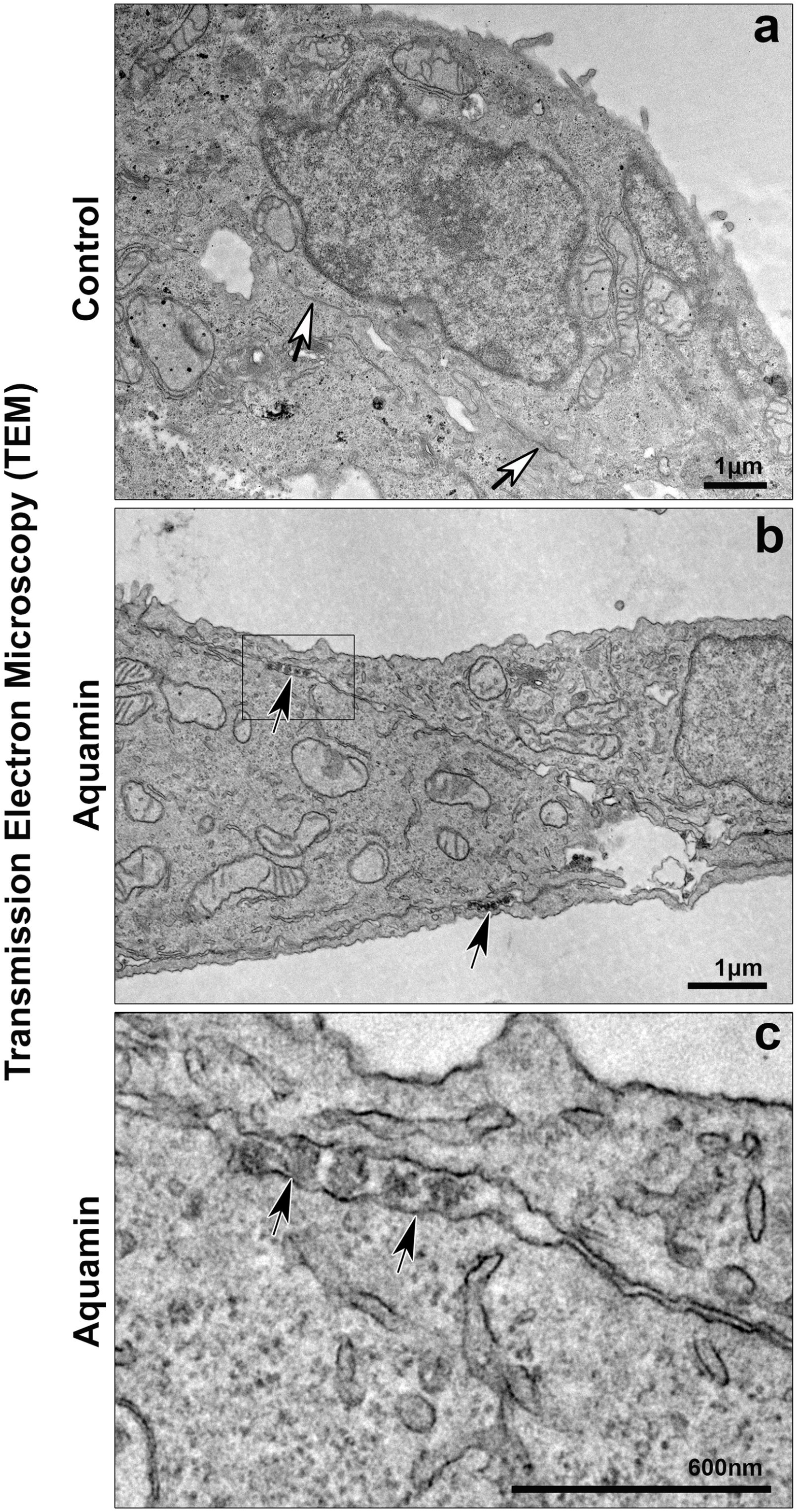
Aquamin-treated Caco-2 cells examined by transmission electron microscopy (TEM). **a**. Control Caco-2 cells. **b** and **c**. Aquamin-treated Caco-2 cells. White arrows in the top panel point to cell-cell border while black arrows in the (panels b and c) point to Aquamin particles in the intercellular space between two adjacent cells. Bars in panels a and b = 1 μm. Bar in panel c = 600 nm.

In addition to utilizing Caco-2 cells as a target for Aquamin binding, we also assessed binding to human colon organoid epithelial cells. Figure 4 demonstrates binding study results with these cells. As can be seen from the series of scanning electron micrographs, binding of particles with a range of size from less than 1 μm to greater than 5 μm in diameter occurred (compare control cells in panel a with Aquamin-treated cells in panels b, c and d). Similar to what was seen with Caco-2 cells, Aquamin binding at cell-cell boundaries was seen with the colon organoid-derived cells (panels e and f, arrows). Careful SEM evaluation of organoid-derived cells that had been exposed to calcium carbonate rarely demonstrated cells with bound material (panel g).

**Figure 4.**
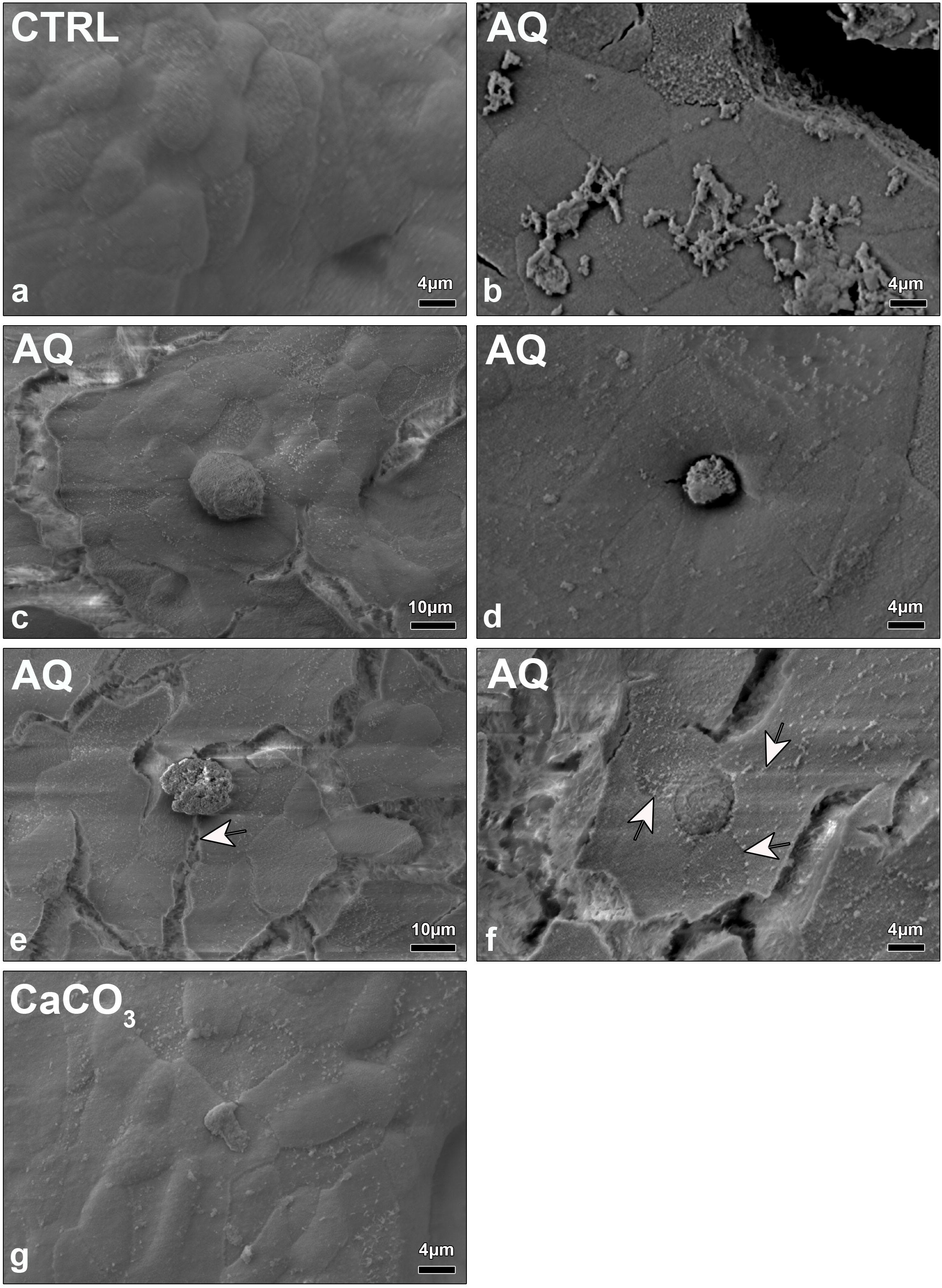
Aquamin binding to human colon organoid epithelial cells. Human colon organoid epithelial cells grown on collagen-IV coated transwell filters were exposed to Aquamin or calcium carbonate as described in the Materials and Methods Section and incubated for 48 hours. At the end of the incubation period, cells were examined by SEM. CTRL = control; AQ = native Aquamin; CaCO_3_ = calcium carbonate. Arrows point to regions of cell-cell contact and associated bound Aquamin particles. Bars in panels a, c, and g = 10 μm; bars in panels b, d, e, and f = 4 μm.

### Effects of Aquamin on expression of proteins involved in cationic element uptake and handling

Based on the data presented above, we conclude that a combination of both soluble and insoluble forms of several important cationic minerals would move through the gastrointestinal tract following Aquamin ingestion. Given this, we asked whether Aquamin, itself, might influence mechanisms responsible for cationic element uptake and handling. To begin addressing this issue, we utilized a proteomic data set generated from a series of human colon organoids (Attili et al. 2019; Aslam et al. 2020b; Varani et al. 2023; Aslam et al. 2025b) to ask whether proteins involved in trace element uptake at the apical surface, intracellular buffering and basolateral export might be responsive to Aquamin. Figure 5 identifies a list of proteins that met the three criteria of i) being recognized in the Reactome database as functionally involved in trace element handling in the gastrointestinal tract, ii) being present in our own colon organoid data set from 12 subjects and iii) being up– or down-regulated by at least 1.2-fold in response to Aquamin. A total of 23 proteins were increased with Aquamin while four were decreased (Figure 5). Among up-regulated proteins were ones that play a direct role in cationic trace element transport across the apical surface of enterocytes (e.g., DMT-1, ZIP4 and CTR1 as well as calcium/manganese antiporter [SLC30A10]). The list of Aquamin-up-regulated proteins also included moieties that function in basolateral export to the circulation – e.g., ceruloplasmin, hephaestin, heme oxygenase-1, peroxiredoxin-6 and CYBRD1. Finally, several other proteins that indirectly influence cationic element transport by facilitating movement of co-transported moieties (e.g., anionic moieties and amino acids) across the gastrointestinal wall as well as moieties that help regulate intracellular pH and ionic strength at the cell surface were also Aquamin-responsive. Among these were two sodium / potassium-transporting ATPases and carbonic anhydrase-4.

**Figure 5.**
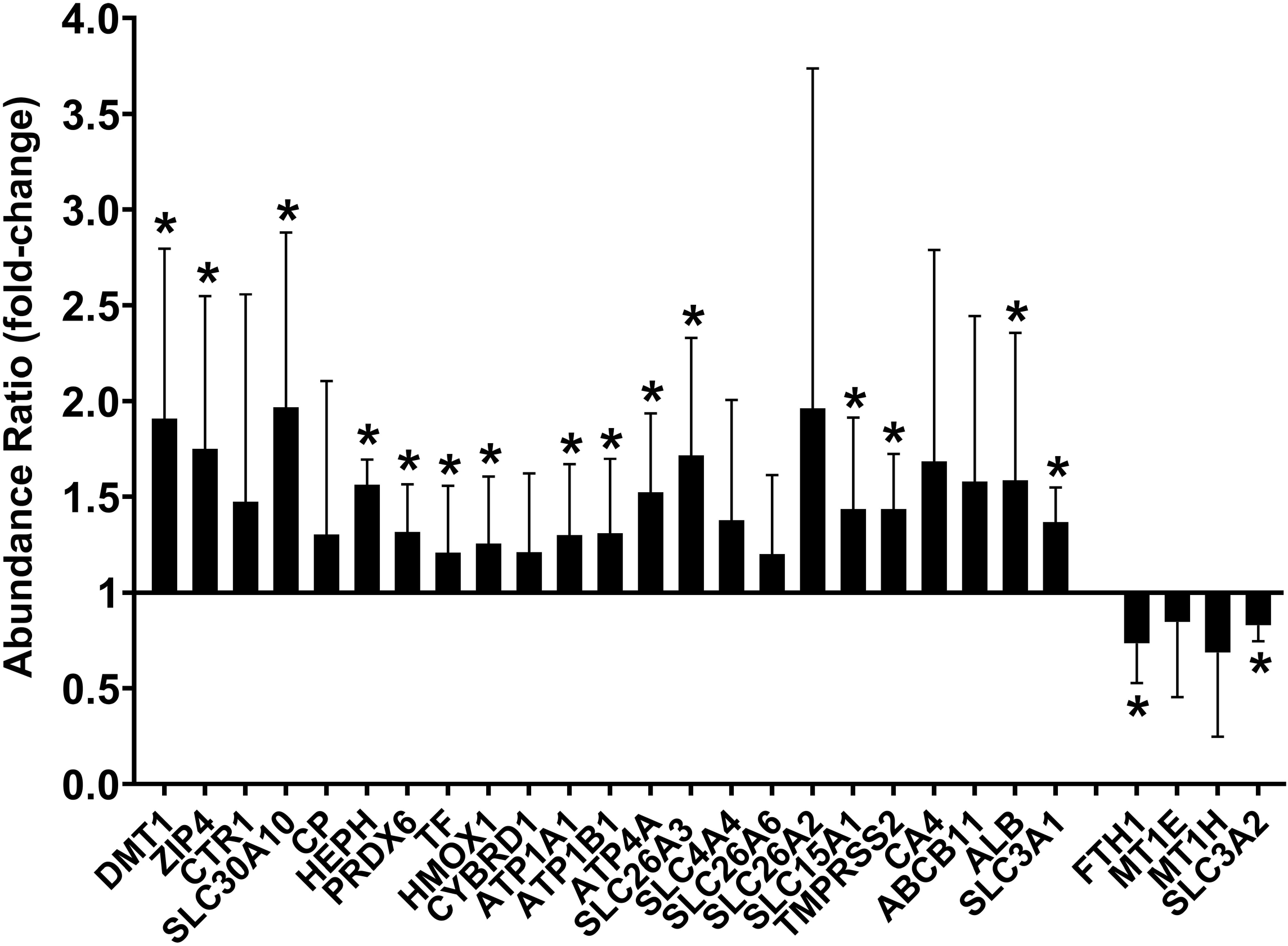
Aquamin effects on proteins involved in cationic element transport in the gastrointestinal tract. Values represent average abundance ratio of Aquamin-treated organoids compared to untreated (control) organoids. Values are means and standard deviations based on merged data from 14 independent proteomic runs with colon organoids from n=12 different subjects. Asterisks (*) indicate statistical difference from control value (i.e., 1.0) at p<0.05 in unpaired t-test.

The down-regulated proteins included metallothionein-1E and –1H, which play a role in intracellular metal ion buffering (Kimura and Kambe 2016), and ferritin. A reduction in the intracellular level of ferritin would “free” iron for export from enterocytes (Camaschella 2019; Rolić et al. 2025; Mahroum et al. 2022; Marques et al. 2022), and its decrease is consistent with up-regulation of hephaestin, peroxiredoxin-6, heme oxygenase-1 and CYBRD1. Together, these findings indicate a coordinated induction of cationic element transport across the gastrointestinal wall with Aquamin. A second (smaller) database generated from (n=3) colon organoids (Attili et al. 2019) treated with calcium alone as well as Aquamin were compared for the same proteins as depicted in Figure 5. Of the 27 proteins in the larger data set, 22 were also present in the smaller set (responsive to Aquamin at ≥1.2-fold change) (Supplement Figure 3). As seen in this figure, the response to Aquamin was similar to what was observed in the larger set (Figure 5). As also seen in Supplement Figure 3, organoid exposure to calcium alone was comparably effective in altering protein levels.

### Trace element analysis in human serum: Effects of daily Aquamin ingestion over a 90 or 180-day period

The protein-profiling data presented above demonstrate that exposure to Aquamin alter*s* expression of several moieties connected to cationic trace element transcellular uptake and handling in the gastrointestinal tract. At the same time, cell binding studies suggest the possibility of paracellular transport. While the data are compelling, neither set of studies proves that trace elements are made available when Aquamin is ingested. As a way to directly determine the effect of Aquamin on cation bioavailability, we utilized serum samples available from two recent clinical trials (Aslam et al. 2020a; Aslam et al. 2021; Aslam et al. 2025a) to assess levels of various trace elements following daily Aquamin ingestion. In the first trial (Aslam et al. 2020a; Aslam et al. 2021), healthy individuals received Aquamin daily for 90 days standardized to provide 800 mg of calcium per day along with the other trace elements in the preparation. In the second trial (Aslam et al. 2025a), subjects with mild UC or UC in remission received the same dose of Aquamin daily for either 90 or 180 days. Supplement Table 4 shows serum levels of the various elements at the pre-treatment and post-treatment visits. As expected, there was little change with most elements since plasma levels of most are tightly regulated (Camaschella 2019; Rolić et al. 2025; López-Alonso et al. 2025; Areco et al. 2020). This was not the case with strontium, however. Increases of 2-3-fold were seen in virtually every study-participant on Aquamin in both trials (Figure 6). Placebo treatment had no effect. Also of interest, in the cohort of UC subjects who ingested Aquamin for 90 to 180 days, serum iron levels increased by 10 and 11% relative to baseline values, respectively, whereas levels decreased by 13% in the placebo group (Supplement Figure 4). Values from healthy subjects did not show this trend. Notably, serum strontium and iron concentrations remained within their respective normal reference ranges.

**Figure 6.**
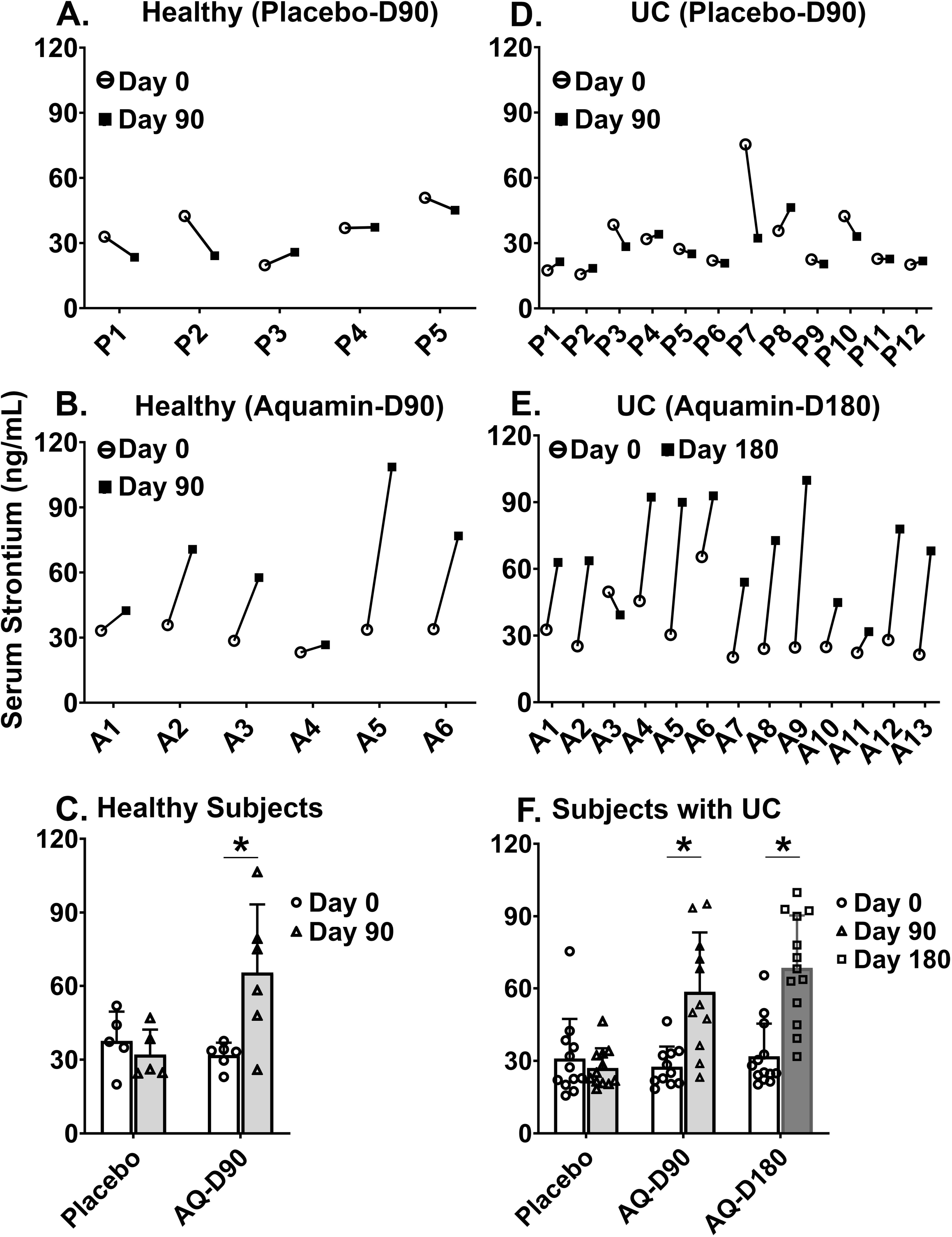
Aquamin effects on strontium levels in human serum. **Upper and middle panels**: Strontium levels in individual serum samples from participants before and after intervention in two recent trials comparing Aquamin groups to placebo. **Lower panels**: Group means and standard deviations. **A-C**: Healthy subjects. **D-E**: Subjects with UC. Statistical analysis was determined using the paired t-test for each set of data separately. Asterisks (*) indicate post-intervention values that are significantly different from pre-intervention value at the p<0.05 level.

## DISCUSSION

Our preclinical studies in rodents (Aslam et al. 2010a; Aslam et al. 2012a; Aslam et al. 2012b; Hampton et al. 2015; Aslam et al. 2010b; Aslam et al. 2013; Aslam et al. 2016; Aviello et al. 2014) and our two interventional trials in human subjects (Aslam et al. 2020a; Aslam et al. 2021; Aslam et al. 2025a; Aslam et al. 2026) have both demonstrated health benefits with Aquamin. In both sets of studies, calcium alone had activity, but the biological response to Aquamin was distinct from and often greater than that seen with calcium alone. Differences between Aquamin and calcium alone may reflect activities of the additional trace elements present in the multi-mineral product. Several of these elements are known to be required for health because deficiencies are associated with overt disease or with deficits in important cellular processes (Camaschella 2019; Rolić et al. 2025; Rosanoff et al. 2012; Prasad 2013; Roohani et al. 2013; Duncan et al. 2015; Melkote et al. 2026; Wang et al. 2023; Rayman 2012; Cardoso et al. 2015; Jiang et al. 2018; Chen et al. 2020; Boivin et al. 1996; Marie et al. 2001). Alternatively, calcium, which is (by far) the most prevalent cationic element in Aquamin (Supplement Table 1) and which plays critical roles in virtually every biological process (Bootman and Bultynck 2020; Stewart et al. 2015; Peterlik et al. 2009), may be largely responsible for the multi-mineral supplement’s activity, with differences in calcium bioavailability between the algae-derived preparation and calcium carbonate being responsible for Aquamin’s enhanced efficacy. The studies described here were carried out to begin addressing this question.

Here we show that Aquamin underwent solubilization in the acid pH (gastric-mimicking) stage-1 of the two-stage digestion simulation protocol. In comparison with calcium carbonate (either limestone or biological materials that are primarily calcium carbonate-based), Aquamin was more acid-resistant as it required more acid and more time to reach the target pH than calcium carbonate. (Supplement Figure 1). When acid-solubilized Aquamin was transferred to the neutral pH environment of stage-2 (mimicking conditions in the small intestine), precipitation occurred. The precipitated material recovered at the end of stage-2 amounted to 15-20% of the starting material by weight. However, when calcium was quantified by ICP-OES, most of the calcium initially present was in soluble form at the end of stage-1 and remained soluble at the end of the stage-2. Findings were similar when calcium carbonate was employed as the starting material. Based on these findings, it is likely that calcium uptake from Aquamin would occur using the same mechanisms employed by calcium from other sources. This would primarily involve transcellular passage of ionic calcium in the small intestine as is well-known (Areco et al. 2020). Paracellular transport (primarily in the small intestine but also occurring in the colon) could also take place, but this would likely constitute a less important route. Paracellular transport (claudin-mediated passage between cells) can involve ionic forms, but calcium-containing nanoparticles have also been shown to be transported through the colonic wall and contribute to the calcium pool under some circumstances (Areco et al. 2020). Our own findings demonstrated binding of Aquamin to the surface of colonic epithelial cells and demonstrated the presence of three-dimensional particles between adjacent cells. Ultimately, there is no *a priori* reason why calcium transport from Aquamin should be distinguishable from other calcium sources in either of these pathways.

Magnesium and strontium were also solubilized during stage-1 digestion and remained mostly soluble through the two-hour period in the stage-2 buffer. Thus, we suggest that these cationic elements would (like calcium) be taken up as ionic forms using known transport pathways. Strontium, because of its similarity to calcium, utilizes the same cellular machinery as calcium for transcellular uptake (though less efficiently) (Bihler et al. 1986; Mathias and Brown 1991; Marcus and Lengemann 1962) and can also be absorbed using paracellular pathways that are employed by calcium. Soluble magnesium can be taken up through both transcellular and paracellular mechanisms, but paracellular absorption plays a relatively greater role in magnesium uptake (Rolić et al. 2025; Liu and Dudley 2025). Thus, even though Aquamin may not be substantially different from other calcium sources in its ability to deliver calcium, its capacity to provide both magnesium and strontium is an important distinction. Both strontium and magnesium have important health benefits (Liu and Dudley 2025; Rosanoff et al. 2012; Jiang et al. 2018; Chen et al. 2020; Boivin et al. 1996; Marie et al. 2001).

Additional cationic trace elements that are known to be important for health (e.g., iron, manganese, copper, zinc, chromium and selenium among them [Camaschella 2019; Rolić et al. 2025; Prasad 2013; Roohani et al. 2013; Duncan et al. 2015; Melkote et al. 2026; Wang et al. 2023; Rayman 2012; Cardoso et al. 2015]) are present in Aquamin. Like calcium, these elements were solubilized from Aquamin in the acid environment of the stage-1 digestion buffer. Unlike calcium, however, these elements were either partially or entirely insoluble by the end of stage-2. Because of this, transcellular passage across the intestinal wall in the small intestine would likely be less efficient than it is with calcium. While the extent to which these important elements are delivered by Aquamin is difficult to assess (due to tight control of plasma levels [Camaschella 2019; Rolić et al. 2025; López-Alonso et al. 2025]), the changes in transporter protein expression observed in colon organoid cultures support the suggestion that Aquamin, itself, promotes uptake of these important trace elements. In regard to iron, specifically, we not only saw increased expression of the major importer protein in enterocytes (i.e., DMT-1) (Camaschella 2019; Rolić et al. 2025) but also elevated expression of several moieties that facilitate iron extrusion from the basolateral surface. At the same time, the level of the intracellular iron-binding protein, ferritin, was down-regulated in Aquamin-treated organoids. Thus, the modest increase in serum iron concentrations observed after Aquamin supplementation in patients with UC is consistent with the suggestion that Aquamin could provide a bioavailable source of this important element. There is no *a priori* reason to believe that other critical elements would not also be available from Aquamin, and no reason to suggest that transcellular mechanisms would not be operative.

Regardless of how effectively most trace elements are provided from Aquamin in ionic forms, delivery of insoluble three-dimensional particles needs to be considered. Trace element delivery as tiny particles via “salvage pathways” in the colon has been documented (Perfecto et al. 2017; Pereira et al. 2013; Domenech et al. 2021; Cubadda et al. 2026). A recent study with iron, specifically, demonstrated endocytic uptake of iron-containing nanoparticles by Caco-2 cells (Pereira et al. 2013), and ultrastructural analysis in the same study suggested dissolution of the particles within the cell. Whether or not Aquamin can deliver a significant amount of one or more cationic trace element via a paracellular pathway is not known. However, evaluation of Aquamin binding to colonic epithelium by both light microscopy and SEM provided evidence that binding occurs readily. SEM revealed that even particles larger than 1 μm in diameter were capable of attaching to the cell surface. SEM also demonstrated particle localization at cell-cell boundaries where paracellular transport mechanisms would be operative. TEM demonstrated the presence of tiny particles in the intercellular space between adjacent cells. Together, these studies suggest that uptake of three-dimensional Aquamin particles is possible. The potential of Aquamin to provide cationic trace elements (including iron) in the insoluble form may be an important consideration since the elements in question have limited solubility in an aqueous environment.

Lanthanide uptake from Aquamin is also of potential interest. While there are known toxicities associated with members of the lanthanide family of rare earth elements, negative consequences reflect pharmacological amounts reaching critical target organs. Lung toxicity has been documented, for example, with inhalation exposure to lanthanum or cerium (Laganà et al. 2026; Kitamura et al. 2007; Valiathan et al. 1986). In the case of gadolinium, intravenous injection of chelated (soluble) forms of this element during MRI-contrast imaging is associated with a rare fibrotic condition. However, this has only been documented with high dose exposure in individuals with compromised kidney function (Cowper et al. 2000; Swartz et al. 2003). Toxicity with any of the lanthanides is not likely to be an issue with Aquamin due to the low abundance (sub-µg/g amounts) of these elements in the natural product and the lack of specific transporters to facilitate transcellular uptake. At low levels, lanthanide elements may actually have a beneficial effect (Li et al. 2021). This is thought to reflect high-affinity binding to the extracellular calcium-sensing receptor (CaSR; Carrillo-López et al. 2010) and concomitant ability to enhance biological responses to calcium (Carrillo-López et al. 2010). The calcium-sensing receptor is not a transporter. It is part of the calcium sensing cascade (Bootman and Bultynck 2020; Stewart et al. 2015). Past studies from our laboratory have demonstrated greater CaSR-induction with Aquamin than with the calcium alone (Singh et al. 2015). The present study demonstrated that these highly-charged trace elements became soluble during stage-1 but rapidly returned to an insoluble state during the second digestion stage. Thus, to the extent that lanthanides are delivered at all from Aquamin, uptake of insoluble three-dimensional particles would likely be the major operative mechanism.

The findings reported here were based, for the most part, on *in vitro* studies. How effectively Aquamin provides a source of cationic elements *in vivo* is not clear. This (lack of clarity) reflects the fact that plasma levels of most elements are tightly controlled, especially when levels are in a physiological concentration range to begin with (Camaschella 2019; Rolić et al. 2025; López-Alonso et al. 2025). Under such conditions, plasma levels do not vary significantly with supplementation, and other relevant tissues are more difficult to access. Of the elements followed in this study, only strontium was consistently elevated in serum after 90 or 180 days of daily Aquamin ingestion compared to pre-intervention levels (Figure 6). Strontium plays a role in metabolic regulation (Jiang et al. 2018; Chen et al. 2020) and, as noted above, is a critical component of bone health (Boivin et al. 1996; Marie et al. 2001). A strontium salt, strontium ranelate, has been developed as a drug for individuals with low bone mass and elevated fracture risk (Marie 2005). Although its efficacy is not in question, the high intake levels of treatment are associated with cardiovascular risks, which have limited its use (Reginster 2014; Atteritano et al. 2016; Donneau and Reginster 2014). Typically, individuals treated with strontium ranelate received high doses (up to 2 grams per day, see Atteritano et al. 2016), resulting in drug concentrations substantially greater than those observed here. Our own studies suggest that benefit from strontium might not require the high drug levels produced with strontium ranelate intervention. For example, our long-term (15-18 month) murine studies demonstrated increased bone mineral density with Aquamin and a concomitant improvement in bone structure / mechanical properties. These changes were associated with a 6-10-fold increase in bone strontium content (Aslam et al. 2010b; Aslam et al. 2013; Aslam et al. 2016). Equally important, individuals in our recent clinical trial demonstrated a statistically significant improvement in bone mineral content and a statistically significant decrease in calculated bone fracture risk identified by DEXA scan imaging before and after 180 days of daily Aquamin ingestion providing only 6 mg of strontium per day (Aslam et al. 2025a). Based on results from our past studies with colon tissue in organoid culture, we have hypothesized that Aquamin’s efficacy primarily reflects local events – i.e., improvement of the colon barrier with a concomitant reduction in inflammation. In support of a direct anti-inflammatory effect, Aquamin has been demonstrated to reduce colitis in a colitis-sensitive mouse model (Aviello et al. 2014) and has been shown to interact with macrophages *in vitro* to interfere with Toll-like receptor signaling (Lioudakis et al. 2026). The findings presented here do not conflict with this past understanding but allow us to suggest that providing a source of critical cationic elements (in addition to calcium) may be another benefit of the multi-mineral product.

These findings need to be tempered by limitations to the present study. Perhaps most important, modeling *in vivo* trace element bioavailability is challenging. Multiple transport pathways (each with multiple proteins) are involved, and efficiency of mineral transport mechanisms vary from individual to individual. In part, these mechanisms are influenced by polymorphisms in critical transport components (da Silva Lopes and Abe 2021; Kiani et al. 2022) but in addition, are altered by age, diet, obesity, physical and psychological stress as well as by chronic inflammatory conditions that affect gut motility and gastrointestinal permeability (López-Alonso et al. 2025). No *in vitro* system can adequately take all of the potential variables into account. Suffice to state is that conditions used here in the two-stage digestion simulation studies (i.e., starting with 800 mg of Aquamin in 20 mL of the stage-1 buffer followed by a 1:2 dilution for stage-2) should be adequate to reflect conditions experienced by participants in our recent clinical studies or those who use Aquamin as an over-the-counter nutritional supplement.

Difficulty in relating cell-binding properties of Aquamin to trace element bioavailability is another limitation. While it is known that sub-micron-sized mineral particles can be taken up by cells of the colon (Pereira et al. 2013), how much this contributes to trace element bioavailability is less clear. Here we have demonstrated that small dispersed Aquamin particles readily attach to the surface of colonic epithelial cells, but what happens after that is not certain. Ultrastructural analysis demonstrated the presence of (sub-micron) Aquamin particles in the intercellular space between adjacent Caco-2 cells, but whether this leads to internalization is difficult to ascertain from static imaging alone. We assume that if internalization does occur, the tiny particles would dissolve in the acid environment within lysosomes, as has been documented with other materials (Pereira et al. 2013). Our data to date are inconclusive and proving this is difficult and warrants additional studies. Whether Aquamin can deliver insoluble trace elements (and to what extent) will require additional studies. Those studies are beyond the scope of the present work.

Finally, our *in vivo* studies convincingly demonstrate strontium delivery but whether other elements are also made available is difficult to prove – again, because serum levels are tightly controlled and other tissue sources that could help address this dilemma are difficult to access in human subjects. All we can say for sure is that the myriads of benefits shown in past studies are unlikely to derive entirely from barrier improvement in the colon. We are currently looking at potential biomarkers that could help predict alterations in the intake of critical trace elements. While previous studies have identified markers for some trace elements, none that are currently available meet all the desired criteria for routine use (López-Alonso et al. 2025). In any event, to address this issue with Aquamin, *per se*, will require new interventional trials and such trials are also beyond the scope of the present work.

In conclusion, deficiencies of multiple trace elements constitute a global public health problem, occurring in both countries with advanced economies and in the developing world (Peterlik et al. 2009; Heaney 2003; Camaschella 2019; Rolić et al. 2025; López-Alonso et al. 2025; Passarelli et al. 2022; Stevens et al. 2022). A low-cost, low-to-no-toxicity natural mineral supplement that could deliver both readily soluble elements as well as trace elements that are difficult to maintain in solution could provide an approach to help mitigate this global problem.

## SUPPLEMENT FIGURES

**Supplement Figure 1**. **Aquamin solubilization in HCl: Comparison with calcium salts.** 100 mg of Aquamin (calcium concentration approximately 32%) was suspended in 2 mL of saline and exposed to 1M HCl in dropwise fashion. The volume of acid needed to bring the powdered material into solution and the time to reach a stable pH of 3.5 were measured. The other calcium-containing materials were obtained from commercial European vendors and treated in the same manner (calcium concentrations approximately 40% except monocalcium phosphate – 17% calcium). Values are based on a single experiment with averages of duplicate samples per data point. The experiment was conducted two times with similar results.

**Supplement Figure 2. Appearance of Aquamin particles after solubilization and precipitation in the two-stage digestion simulation: Comparison with native Aquamin and calcium carbonate**. Bars in a, c, and e = 20 µm. Bars in b, d, and f = 4 µm.

**Supplement Figure 3. Aquamin effects on proteins involved in cationic element transport in the gastrointestinal tract: Comparison with calcium alone**. Values represent average abundance ratio of Aquamin-treated organoids compared to untreated (control) organoids and the average abundance ratio of calcium-treated organoids compared to untreated (control) organoids. Values are means and standard deviations based on merged data from 3 independent proteomic runs comparing Aquamin to calcium alone in each proteomic run. Asterisks (*) indicate statistical difference from control value (i.e., 1.0) at p<0.05 in unpaired t-test.

**Supplement Figure 4. Aquamin effects on iron levels in human serum**. **Upper and middle panels**: Iron levels in individual serum samples from participants before and after intervention in two recent trials. **Lower panels**: Group means and standard deviations. **A-C**: Healthy subjects. **D-E**: Subjects with UC.

## SUPPLEMENT TABLES

Supplement Table 1. Distribution of 12 cationic trace elements in Aquamin

Supplement Table 2. Solubility of 12 cationic trace elements as a function of time in the stage-2 digestion buffer

Supplement Table 3. Calcium carbonate solubilization in the two-stage digestion simulation

Supplement Table 4. Serum levels of 12 cationic trace elements

## SUPPLEMENT TEXT FILES

Supplementary File 1. Methods related to “Inductively Coupled Plasma (ICP) Analysis (ICP-OES and ICP-MS).”

## Acknowledgments.

Human colon organoid-derived epithelial cells used in three-dimensional (3D) cultures to assess mineral binding were obtained from the University of Michigan Human Organoid Core. Scanning electron microscopy (SEM) and transmission electron microscopy (TEM) analyses were conducted in collaboration with the Microscopic Imaging Core Laboratory at the University of Michigan. Elemental analyses of samples from the digestion studies and serum samples were performed at the Michigan State University Quantitative Bio Element Analysis and Mapping (QBEAM) Center. We also thank the Proteomics Resource Facility at the University of Michigan for assistance with proteomic data acquisition. We are grateful to Marigot Ltd. (Cork, Ireland) for providing Aquamin® as a gift. Most importantly, we thank the study participants, including healthy volunteers and patients with ulcerative colitis, without whom this research would not have been possible.

## Data availability statement

The datasets generated and analyzed in this study are available through an online repository. The mass spectrometry proteomics data have been deposited in the ProteomeXchange Consortium via the PRIDE partner repository; the dataset identifier is pending.

## Ethics statement

For the 3D culture study, de-identified colon tissue was obtained during routine colonoscopies from healthy participants or from deceased donors through the Gift of Life program. Tissue collection was conducted under a protocol reviewed and approved by the University of Michigan Medical School Institutional Review Board (IRBMED). No protected health information was shared with the investigators. Participants provided written informed consent before the procedure, authorizing the use of their colonic tissue for research purposes. Serum samples used in this study were obtained from participants enrolled in the clinical trials conducted by Aslam et al. The clinical trial was conducted under IRBMED oversight and an investigational new drug (IND) application approved by the U.S. Food and Drug Administration (FDA).

