## Supplementary figures and images for "Aquamin: A Source of Bioavailable Calcium, Magnesium and Additional Cationic Trace Elements in Colon Health and Beyond"

### Supplement Figure 1

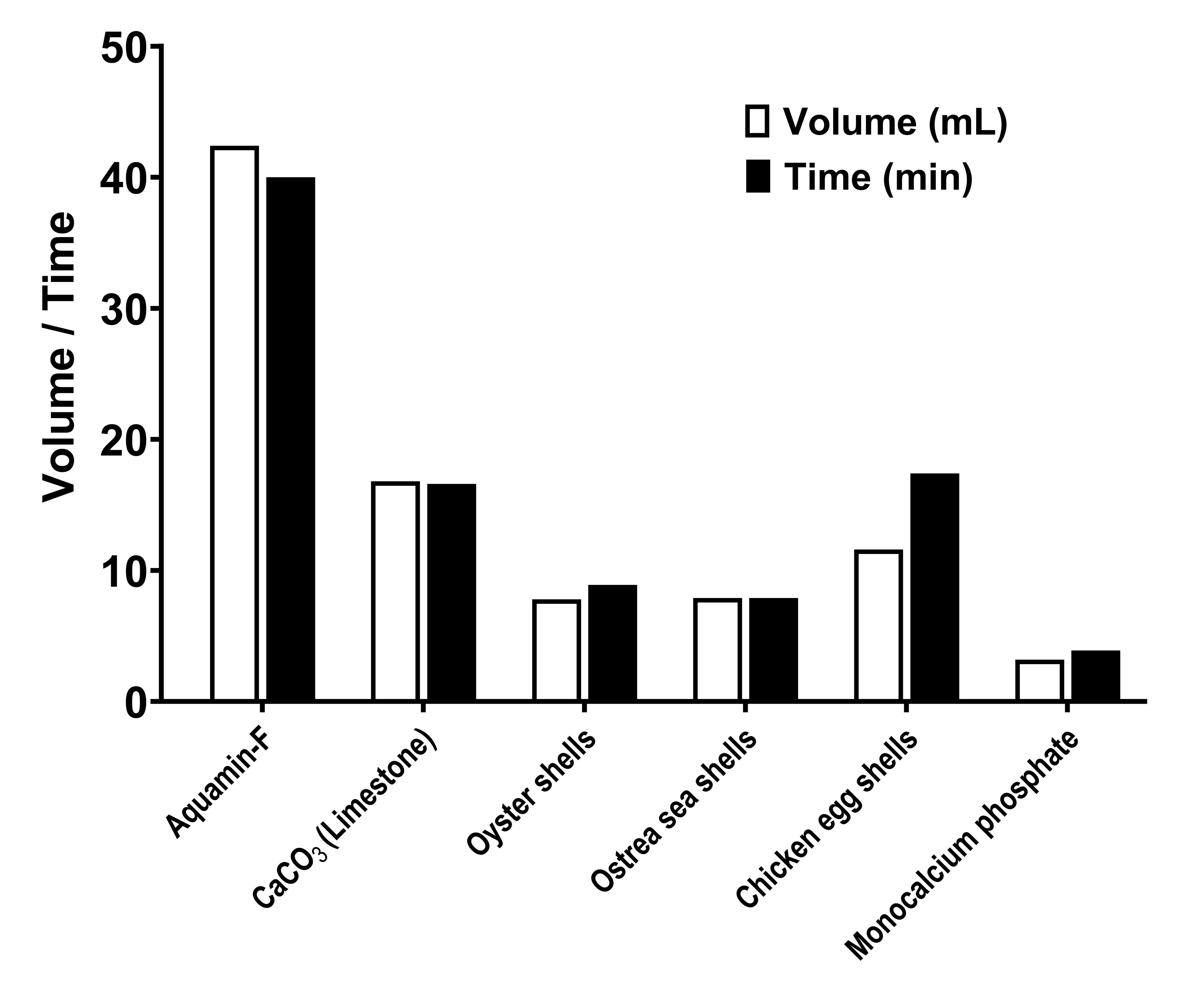

### Supplement Figure 2

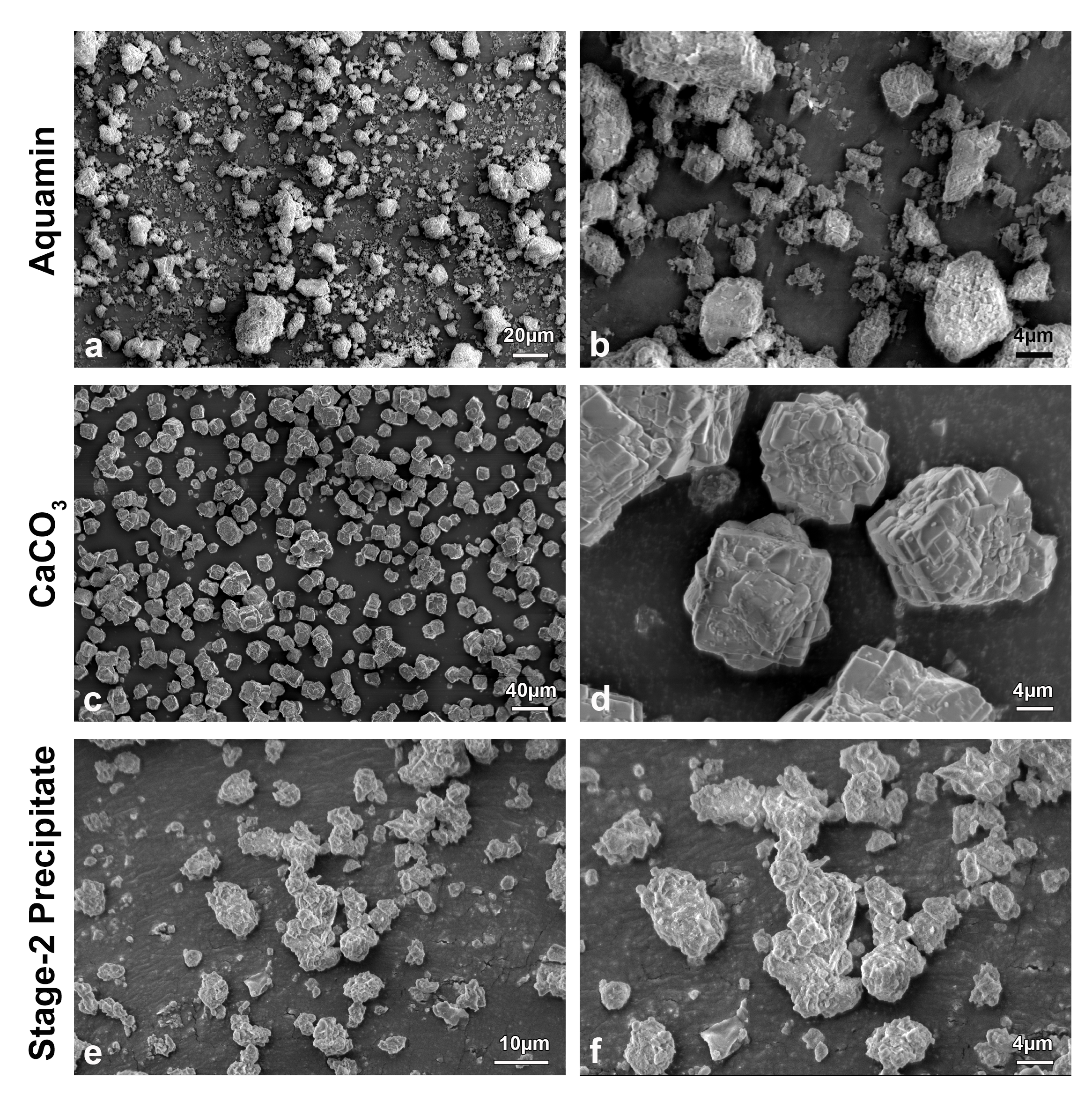

### Supplement Figure 3

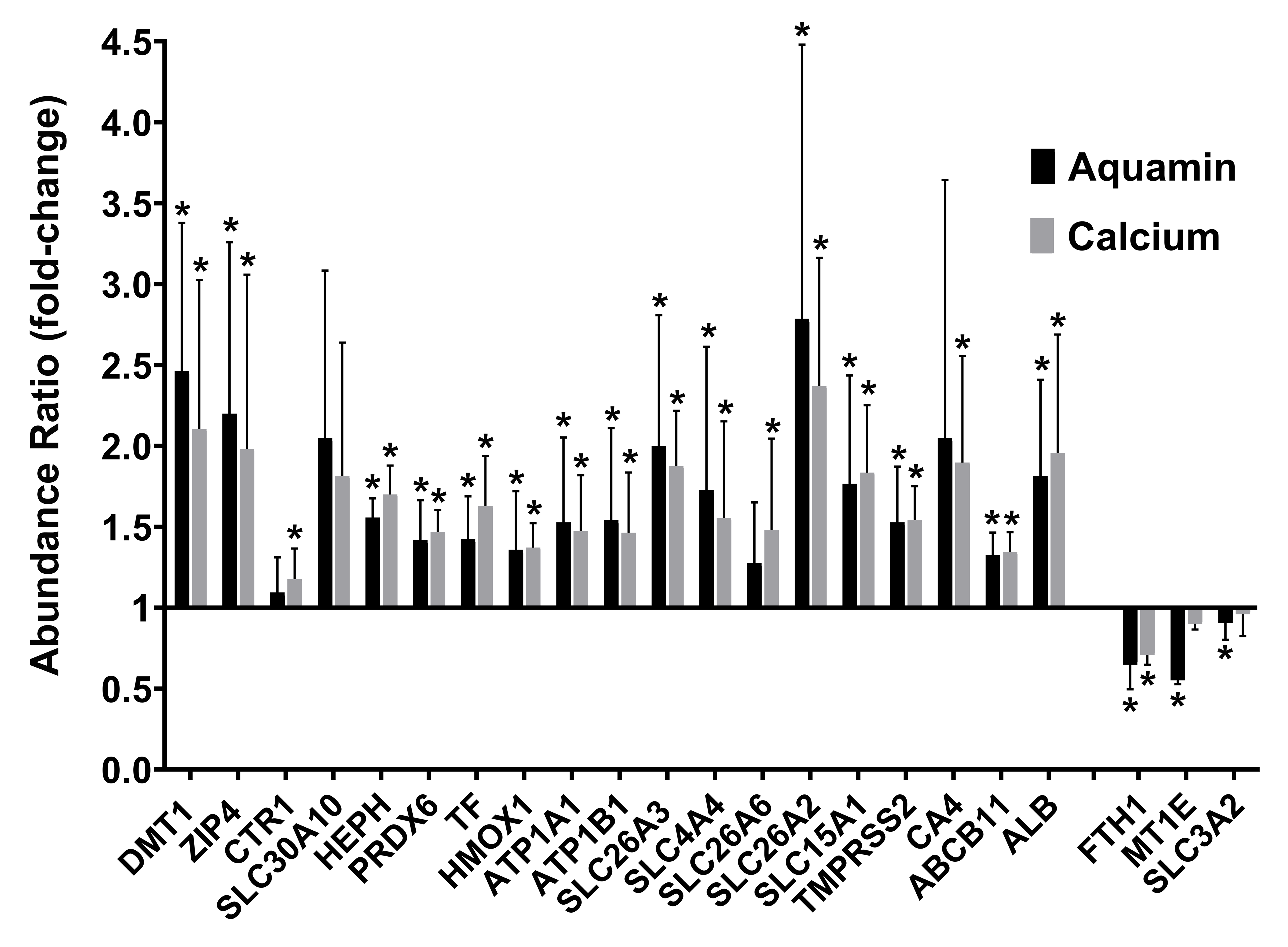

### Supplement Figure 4

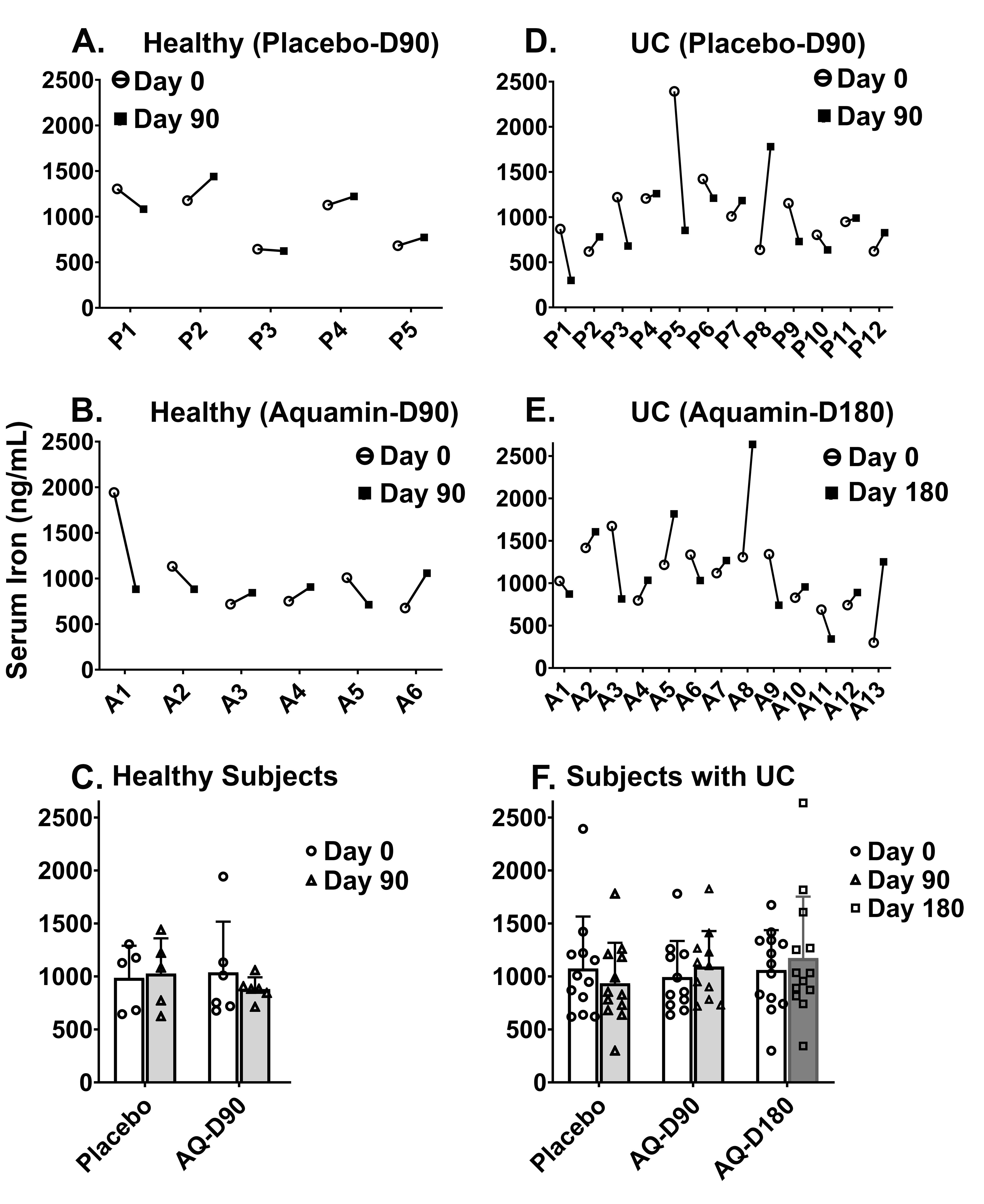
