## Supplement File 1 for "Aquamin: A Source of Bioavailable Calcium, Magnesium and Additional Cationic Trace Elements in Colon Health and Beyond"

**Supplementary File 1.**

**Inductively Coupled Plasma (ICP) Analysis (ICP-OES and ICP-MS).** Elemental analysis was performed at the Michigan State University Quantitative Bio Element Analysis and Mapping (QBEAM) Center. All standards, blanks, and samples were prepared using sub-boiling point distilled (Savillex DST-1000, Eden Prairie, MN, USA) trace metal grade nitric acid (70%, Fisher chemical, Cat# A509P212), ultrapure water (18.2 MΩ·cm @ 25°C), and metal-free polypropylene conical tubes (15 and 50 mL, Labcon, Petaluma, CA, USA) and measured gravimetrically. Samples for ICP-OES and ICP-MS were prepared in an identical manner. Specifically, samples generated in the two-stage digestion simulation protocol were diluted 1:10 with 3% nitric acid. Serum samples were diluted 1:100 with 3% nitric acid. Both sample types were then incubated at room temperature for at least 1 hour resulting in complete solubilization. Standards were prepared with standard IV-ICPMS-71A (Inorganic Ventures; Christiansburg, VA). A high standard of 100 ppb with 1:2 serial dilutions down to 6.25 ppb in 3% (v/v) trace nitric acid in ultrapure deionized water was used to generate standard curves.

***ICP-OES Analysis***

ICP-OES samples, standards, and blanks were analyzed using an Agilent 5800 ICP-OES (Agilent, Santa Clara, CA, USA) equipped with the Agilent SPS 4 Autosampler, quartz cyclonic spray chamber, micromist nebulizer, and AVS 6/7 valve introduction system. Daily instrument performance was validated via a detector calibration and multiwavelength calibration in axial and radial mode using the manufacturer’s multi-element calibration standard containing 50 mg/L of the following elements: Al, As, Ba, Cd, Co, Cr, Cu, Mn, Mo, Ni, Pb, Se, Sr, Zn and 500 mg/L K in 5% nitric acid. Internal standardization was accomplished inline using the AVS 6/7 valve and a 1 µg/g Y internal standard solution (Inorganic Ventures, Christiansburg, VA, USA) in 3% (v/v) trace nitric acid in ultrapure water.

System parameters were as follows: Read time = 5 s (3 replicates), RF power = 1.2 kW, Stabilization time = 5 s, nebulizer flow = 0.70 L/min, plasma flow = 12.0 L/min, Aux flow = 1.00 L/min, Make up flow = 0.00 L/min, and Radial Viewing height = 8 mm. Wavelengths selected for analysis were all in Radial mode and were Ca (422.673 nm), K (766.491 nm), Mg (285.213 nm), Na (589.592 nm), P (213.618 nm), S (181.972 nm) using Y (371.029 nm) as the internal standard. Standards were prepared at 100, 50, 25, 12.5, 6.25, 3.125, 1.5625, and 0 µg/g multi-element standard and fit using linear regression with no weighting with an R^2^ > 0.999 and %RSE < 10% . Sample sequence was run with a continuing calibration blank (CCB) every 3 samples and continuing calibration verification (CCV) every 30 samples.

***ICP-QQQ-MS Analysis***

ICP-MS samples, standards, and blanks were analyzed using an Agilent 8900 Triple Quadrupole ICP-MS (Agilent, Santa Clara, CA, USA) equipped with the Agilent SPS 4 Autosampler, integrated sample introduction system (ISiS), x-lens, and Micromist nebulizer. Daily tuning of the instrument was accomplished using the manufacturer supplied tuning solution containing Li, Co, Y, Ce, and Tl. Global tune optimization was based on optimizing intensities for ^7^Li, ^89^Y, and ^205^Tl while minimizing oxides (^140^Ce^16^O/^140^Ce < 1.5%) and doubly charged species (^140^Ce++/^140^Ce+ < 2%). Following global instrument tuning, gas mode tuning in He KED was accomplished using the same manufacturer supplied tuning solution. In KED mode (using 100% UHP He, Airgas), intensities for ^59^Co, ^89^Y, and ^205^Tl were maximized while minimizing oxides (^140^Ce^16^O/^140^Ce < 0.5%) and doubly charged species (^140^Ce++/^140^Ce+ < 1.0%) with short-term RSDs < 3.5%. Internal standardization was accomplished inline using the ISIS valve and a 200 ng/g of mixed single element standards multi-element internal standard mix containing Bi, In, ^6^Li, Sc, Tb, Y (IV-ICPMS-71D, Inorganic Ventures, Christiansburg, VA, USA) in 3% (v/v) trace nitric acid in ultrapure water. Acquisition parameters were as follows: gas mode = He KED, Acquisition mode = Spectrum, Replicates = 3, Sweeps/replicate = 30, Stabilization time = 10 s, Scan type = Single quad, RF power = 1550 W, Sampling Depth = 8 mm, Nebulizer gas = 1.09 L/min, Plasma gas = 15 L/min, Auxiliary gas = 0.9 L/min, He flow rate = 5 mL/min, OctP Bias = -18.0 V, Axial acceleration = 1 V, Energy discrimination = 3.0 V. Isotopes selected for analysis were: ^44^Ca, ^52^Cr, ^55^Mn, ^56^Fe, ^59^Co, ^60^Ni, ^65^Cu, ^66^Zn, ^82^Se, ^88^Sr, ^139^La, ^140^Ce, ^157^Gd with ^45^Sc, ^89^Y, ^115^In, ^159^Tb, and ^209^Bi selected as internal standards. Standards were prepared at 100, 50, 25, 12.5, 6.25, and 0 ng/g multi-element standard and fit using linear regression with no weighting with an R^2^ > 0.9999 and %RSE < 5%. Sample sequence was run with a continuing calibration blank (CCB) every 5 samples and continuing calibration verification (CCV) every 30 samples.

**Abbreviations.**

Symbol	Element

Al	Aluminum

As	Arsenic

Ba	Barium

Bi	Bismuth

Ca	Calcium

Cd	Cadmium

Ce	Cerium

Co	Cobalt

Cr	Chromium

Cu	Copper

Fe	Iron

Gd	Gadolinium

In	Indium

K	Potassium

La	Lanthanum

Li	Lithium

Mg	Magnesium

Symbol	Element

Mn	Manganese

Mo	Molybdenum

Na	Sodium

Ni	Nickel

O	Oxygen

P	Phosphorus

Pb	Lead

S	Sulfur

Sc	Scandium

Se	Selenium

Sr	Strontium

Tb	Terbium

Tl	Thallium

Y	Yttrium

Zn	Zinc
