## Supplement Table 1 for "Aquamin: A Source of Bioavailable Calcium, Magnesium and Additional Cationic Trace Elements in Colon Health and Beyond"

**Supplement Table 1. Distribution of 12 cationic trace elements in Aquamin**

Trace element	Amount in Aquamin (mg/kg)

(mean ± SD)	____________________________________________________________________________

| Calcium | 321,325 ± 26,216 |
| --- | --- |
| Magnesium | 26,226 ± 1975 |
| Strontium | 2371 ± 202 |
| Iron | 750 ± 95 |
| Manganese | 62 ± 16 |
| Copper | 4.16 ± 1.0 |
| Zinc | 11.95 ± 4.4 |
| Chromium | 2.00 ± 0.09 |
| Selenium | 1.19 ± 0.05 |
| Lanthanum | 0.67 ± 0.11 |
| Cerium | 1.34 ± 0.12 |
| Gadolinium | 0.27 ± 0.09 |

__________________________________________________________________________

ICP-OES and ICP-MS was used to assess levels of all detectable elements in four separate lots of Aquamin. Values shown are means and standard deviations of the top 12 elements of interest.
