## Supplement Table 2 for "Aquamin: A Source of Bioavailable Calcium, Magnesium and Additional Cationic Trace Elements in Colon Health and Beyond"

**Supplement Table 2. Solubility of 12 cationic trace elements as a function of time in the stage-2 digestion buffer**

| Trace element | Percent soluble | | | | | | |
| --- | --- | --- | --- | --- | --- | --- | --- |
|  | 5 min | 10 min | 15 min | 30 min | 60 min | 90 min | 120 min |
| Calcium | >99 | 98 | >99 | 98 | >99 | 98 | 96 |
| Magnesium | >99 | 98 | >99 | 98 | >99 | >99 | >99 |
| Strontium | >99 | 95 | 99 | 94 | >99 | 98 | 97 |
| Iron | 3.2 | 2.3 | 1.8 | 0.77 | 0.27 | 0.24 | 0.25 |
| Manganese | 82 | 79 | 82 | 77 | 79 | 78 | 77 |
| Copper | 43 | 41 | 43 | 42 | 44 | 42 | 49 |
| Zinc | 21 | 14 | 16 | 15 | 14 | 18 | 11 |
| Chromium | 4.2 | 3.9 | 3.9 | 3.7 | 4.7 | 2.8 | 5.4 |
| Selenium | 45 | 50 | 29 | 38 | 60 | 45 | 64 |
| Lanthanum | <0.10 | <0.10 | <0.10 | <0.10 | <0.10 | <0.10 | <0.10 |
| Cerium | <0.10 | <0.10 | <0.10 | <0.10 | <0.10 | <0.10 | <0.10 |
| Gadolinium | <0.10 | <0.10 | <0.10 | <0.10 | <0.10 | <0.10 | <0.10 |

The two-stage digestion simulation was carried out. Duplicate samples were obtained from the stage-2 digestion buffer at times between 5 minutes and 120 minutes. One sample was assessed “as is” while the other sample was filtered through a 0.22 μm filter and then assessed. ICP-OES and ICP-MS were used to assess levels of all detectable elements in each sample. Percent soluble values were determined by comparing values in filtered samples to unfiltered samples. Values shown represent the percent soluble at each timepoint.
