## Supplement Table 3 for "Aquamin: A Source of Bioavailable Calcium, Magnesium and Additional Cationic Trace Elements in Colon Health and Beyond"

**Supplement Table 3. Calcium carbonate solubilization in the two-stage digestion simulation**

Sample	Calcium level	% Soluble

(ICP-OES value)

______________________________________________________________________________

Stage-1 total	13315 ± 730

Stage-1 filtered	13055 ± 762	98 ± 1

Stage-2 total	12670 ± 290

Stage-2 filtered	12650 ± 594	98 ± 2

________________________________________________________________________________

The two-stage digestion simulation was carried out with calcium carbonate as the starting material. Duplicate samples were obtained from the stage-1 and stage-2 digestion buffers at the end of the incubation period. One sample from each stage was assessed “as is” while the other sample was filtered through a 0.22 μm filter and then assessed. ICP-OES was used to assess calcium levels in each sample (presented in µg/g). Percent soluble values were determined by comparing values in filtered samples to unfiltered samples.
