## Supplement Table 4 for "Aquamin: A Source of Bioavailable Calcium, Magnesium and Additional Cationic Trace Elements in Colon Health and Beyond"

| **Supplement Table 4. Serum levels of 12 cationic trace elements** | | | | | | | | |
| --- | --- | --- | --- | --- | --- | --- | --- | --- |
|  | *Healthy Subjects* | | | | |  |  |  |
| Element (ng/mL) | AQ-D90 | |  | Placebo | |  |  |  |
|  | Day 0 | Day 90 |  | Day 0 | Day 90 |  |  |  |
| Calcium | 84610±4513 | 91505±7968 |  | 100894±25973 | 92750±8981 |  |  |  |
| Magnesium | 18852±1370 | 20579±1763 |  | 19456±3016 | 18785±1674 |  |  |  |
| Strontium | 31.88±5.03 | 65.44±27.87 |  | 37.64±11.89 | 32.08±10.14 |  |  |  |
| Iron | 1039±478 | 881±111 |  | 987±303 | 1028±331 |  |  |  |
| Chromium | 0.27±0.00 | ND |  | 0.20±0.00 | 7.33±0.00 |  |  |  |
| Manganese | 2.69±0.81 | 3.37±1.47 |  | 5.18±1.60 | 5.10±1.52 |  |  |  |
| Copper | 1073±187 | 1126±256 |  | 1294±277 | 1185±230 |  |  |  |
| Zinc | 775±103 | 793±149 |  | 792±102 | 768±43.41 |  |  |  |
| Selenium | 30.81±13.50 | 28.83±3.81 |  | 42.19±20.38 | 35.34±6.81 |  |  |  |
| Lanthanum | 0.06±0.10 | 0.03±0.02 |  | 0.03±0.02 | 0.02±0.01 |  |  |  |
| Cerium | 4.75±0.70 | 4.99±1.16 |  | 5.59±0.98 | 5.09±0.80 |  |  |  |
| Gadolinium | 0.13±0.22 | 0.06±0.08 |  | 0.02±0.01 | 0.02±0.00 |  |  |  |
|  | *Subjects with UC* | | | | | | | |
| Element (ng/mL) | AQ-D180 | |  | AQ-D90 | |  | Placebo | |
|  | Day 0 | Day 180 |  | Day 90 | Day 180 |  | Day 0 | Day 90 |
| Calcium | 86458±14060 | 86228±15024 | | 97512±15716 | 105428±28591 | | 100760±14255 | 97512±15716 |
| Magnesium | 19342±2827 | 19422±1802 |  | 18487±2978 | 18623±3542 |  | 18832±2543 | 18487±2978 |
| Strontium | 32.76±13.72 | 68.51±22.83 |  | 27.07±8.13 | 59.39±23.57 |  | 30.98±16.36 | 27.07±8.13 |
| Iron | 1061±375 | 1175±578 |  | 994±340 | 1094±334 |  | 1075±490 | 936±382 |
| Chromium | 1.48±2.05 | 0.68±0.54 |  | 0.66±0.30 | 0.86±1.37 |  | 1.11±1.08 | 0.66±0.30 |
| Manganese | 6.75±5.76 | 5.01±3.18 |  | 3.67±2.81 | 3.38±2.67 |  | 3.77±2.72 | 3.67±2.81 |
| Copper | 1151±537 | 1063±419 |  | 1038±310 | 995±258 |  | 1069±251 | 1038±310 |
| Zinc | 772±160 | 776±136 |  | 785±164 | 779±190 |  | 802±72.17 | 785±164 |
| Selenium | 51.26±21.91 | 48.47±18.89 |  | 47.51±17.95 | 48.23±17.44 |  | 49.96±20.14 | 47.51±17.95 |
| Lanthanum | 0.09±0.08 | 0.11±0.13 |  | 0.54±1.62 | 0.06±0.05 |  | 0.12±0.19 | 0.54±1.62 |
| Cerium | 4.74±1.85 | 5.44±3.42 |  | 5.65±5.38 | 4.41±1.28 |  | 4.58±0.78 | 5.65±5.38 |
| Gadolinium | 0.19±0.51 | 0.05±0.06 |  | 0.02±0.02 | 0.01±0.01 |  | 0.03±0.03 | 0.02±0.02 |

*ND: Not detected.

Serum samples were collected from the subjects before and after the intervention as part of the studies. The study involving healthy subjects (n = 5-6) lasted 90 days, whereas the study involving subjects with ulcerative colitis (UC; n = 11-13) lasted up to 180 days.
